# Depletion of hemoglobin transcripts and long read sequencing improves the transcriptome annotation of the polar bear (*Ursus maritimus*)

**DOI:** 10.1101/527978

**Authors:** Ashley Byrne, Megan A. Supple, Roger Volden, Kristin L. Laidre, Beth Shapiro, Christopher Vollmers

## Abstract

Transcriptome studies evaluating whole blood and tissues are often confounded by overrepresentation of highly abundant transcripts. These abundant transcripts are problematic as they compete with and prevent the detection of rare RNA transcripts, obscuring their biological importance. This issue is more pronounced when using long-read sequencing technologies for isoform-level transcriptome analysis, as they have relatively lower throughput compared to short-read sequencers. As a result long-read based transcriptome analysis is prohibitively expensive for non-model organisms. While there are off-the-shelf kits available for select model organisms capable of depleting highly abundant transcripts for alpha (HBA) and beta (HBB) hemoglobin, they are unsuitable for non-model organisms. To address this, we have adapted the recent CRISPR/Cas9 based depletion method (Depletion of Abundant Sequences by Hybridization) for long-read full-length cDNA sequencing approaches that we call Long-DASH. Using a recombinant Cas9 protein with appropriate guide RNAs, full-length hemoglobin transcripts can be depleted in-vitro prior to performing any short- and long-read sequencing library preparations. Using this method, we sequenced depleted full-length cDNA in parallel using both our Oxford Nanopore Technology (ONT) based R2C2 long-read approach, as well as the Illumina short-read based Smart-seq2 approach. To showcase this, we have applied our methods to create an isoform-level transcriptome from whole blood samples derived from three polar bears (*Ursus maritimus*). Using Long-DASH, we succeeded in depleting hemoglobin transcripts and generated deep Smart-seq2 Illumina datasets and 3.8 million R2C2 full-length cDNA consensus reads. Applying Long-DASH with our isoform identification pipeline, Mandalorion we discovered ~6,000 high-confidence isoforms and a number of novel genes. This indicates there is a high diversity of gene isoforms within Ursus maritimus not yet reported. This reproducible and straightforward approach has not only improved the polar bear transcriptome annotations but will serve as the foundation for future efforts to investigate transcriptional dynamics within the 19 polar bear subpopulations around the Arctic.

## Introduction

Accurate isoform-level differential expression analysis of transcriptomes is essential for interpreting gene regulation under different biological, environmental or physiological conditions. RNA transcript isoforms – which are often unique among different cell types, tissues, developmental stages, and organisms (1–3) – are defined by the use of alternative transcription start sites (TSSs), polyA sites, and splice sites. Use of alternative isoforms is highly regulated and thought to contribute to cellular and organismal diversification within higher eukaryotes (4), adaptation and speciation (5, 6) and can also reflect certain disease states (7–9).

To perform this type of differential expression analysis on the isoform-level requires both short- and long-read sequencing technology. Short-read RNA-seq provides the read depth necessary for gene expression quantification but requires accurate and exhaustive isoform-level transcriptome annotations for its analysis. However, existing transcriptome annotations of non-model organisms are often incomplete or inaccurate (10) because they cannot rely on labor-intensive efforts like Gencode, which are working to exhaustively annotate the isoform-level transcriptomes of human and mouse. While short-read RNA-seq data can itself be used for transcriptome annotation, it fails at annotating transcriptomes on the isoform-level because it cannot recapitulate full-length transcripts. This inability to define full-length transcripts is due to the fragmentation of RNA, or their cDNA copies, prior to sequencing making it difficult to computationally re-assemble reliably (11–13). To provide an accurate isoform-level transcriptome annotation for non-model organisms, long-read sequencing technology is required to sequence full-length cDNA molecules.

The ability to perform combined short- and long-read transcriptome analysis on non-model organisms is further complicated by sample availability. In contrast to the organs and tissues of model organisms which can be easily acquired, availability of samples from non-model organisms are often more limited. In rare circumstances sampling can be performed through fat and muscle tissue biopsies (14), but the current gold standard still relies on whole blood RNA samples, especially for large non-model organisms (15). This is particularly true for protected and endangered species (16, 17). While whole blood samples can be easily acquired and provide a wealth of information regarding physiological or disease states in surrounding tissues (18), polyadenylated RNA extracted from whole blood can be comprised of >50% hemoglobin transcripts (19, 20). In any high-throughput sequencing-based assay, these highly abundant transcripts will compete for a limited number of sequencing reads and as a result will be sequenced over and over again without generating any new information. This would waste valuable reads which could otherwise detect less abundant transcripts.

Currently, there is no approach to deplete hemoglobin transcripts from whole blood RNA while enabling downstream analysis of the depleted RNA/cDNA with both short- and long-read sequencing. Commercially available hemoglobin depletion kits – including GLOBINclear (Ambion) or RiboZero (Illumina) – are specifically designed for human samples and rely on hemoglobin RNA pull-down methods (21). Even if they would succeed in depleting hemoglobin from non-model organism samples, which is far from guaranteed (22), these pull-down approaches use harsh conditions and high temperatures during long incubation steps which contribute to RNA fragmentation and introduce unwanted technical variability (23). While fragmented RNA is suitable as input into short-read RNA-seq, it is not suitable for long-read full-length cDNA sequencing.

To perform a comprehensive analysis of non-model organism transcriptomes from whole-blood with short- and long-read technologies, we require a new approach that can deplete highly abundant transcripts like hemoglobin from whole-blood samples of a wide range of organisms without fragmenting transcripts. To this end, we chose to adapt the powerful, recently published DASH (Depletion of Abundant Sequences by Hybridization) (24) method which utilizes a recombinant Cas9 to perform in-vitro depletion using sequence specific sgRNAs. Our adapted method which we will refer to as Long-DASH also takes advantage of the CRISPR/Cas9 system to selectively deplete hemoglobin alpha (HBA) and beta (HBB) transcripts but targets full-length cDNA instead of fragmented short-read Illumina sequencing libraries like regular DASH. By depleting full-length cDNA prior to any library preparation, this allows the user the choice to use any short- or long-read sequencing platform.

As a proof-of-concept we evaluated three hemoglobin-depleted and non-depleted polar bear whole blood transcriptomes using our ONT-based R2C2 (25) full-length cDNA sequencing method and an Illumina-based modified Smart-seq2 method. We found that by incorporating Long-DASH, we successfully depleted hemoglobin transcripts without non-specifically affecting the rest of the cDNA pool. Finally, we generated ~3.8 million ONT-based R2C2 consensus reads, dramatically refining the polar bear transcriptome annotations.

## Results

### Long-DASH depletes hemoglobin transcripts from full-length cDNA

We used a modified Smart-seq2 protocol (25–27) to reverse transcribe and amplify full-length cDNA from 70 ng of whole blood RNA of three polar bears (PB3, PB19, PB21). We then performed a targeted depletion of hemoglobin transcripts by incubating the full-length cDNA with Cas9 protein loaded with 16 guide RNAs (sgRNAs) specific to hemoglobin transcripts – 8 sgRNAs targeting the HBA transcripts and 8 sgRNAs targeting the HBB transcripts. The sgRNAs were selected to deplete hemoglobin transcripts from human and polar bear samples. The sgRNAs were chosen based upon sequence homology between these two species to eventually allow the removal of abundant of hemoglobin transcripts in whole blood from both human and polar bear samples using the same sgRNAs (21) (Supplementary Fig.1). Using the 16 sgRNA probes we designed should also allow for the depletion of samples of other vertebrates although sequence similarity should be checked before this is attempted.

The depletion process using the Cas9 system should cut the ~700-800 bp transcripts at different sites allowing us to do two things. First, we can re-amplify the sample, thereby only enriching for full-length molecules since the cut cDNA molecules no longer contain two priming sites required for exponential amplification during PCR amplification (Fig. 1). Second, we can remove the cut transcripts by performing a SPRI-bead based size selection whereby only transcripts > 500 bp are retained. Indeed, prior to any depletion, we observed very strong bands located at ~700-800 bp in our agarose gels indicating the presence of a substantial amount of HBA and HBB hemoglobin transcripts (Fig. 2). After depletion, reamplification, and size selection, the full-length cDNA product was visualized again to reveal the removal of the putative hemoglobin bands (Fig. 2). After hemoglobin depletion is confirmed, the cDNA is ready to be converted into ONT- and Illumina-based libraries, with each protocol using the same input cDNA (Fig. 1).

**Figure 1:**
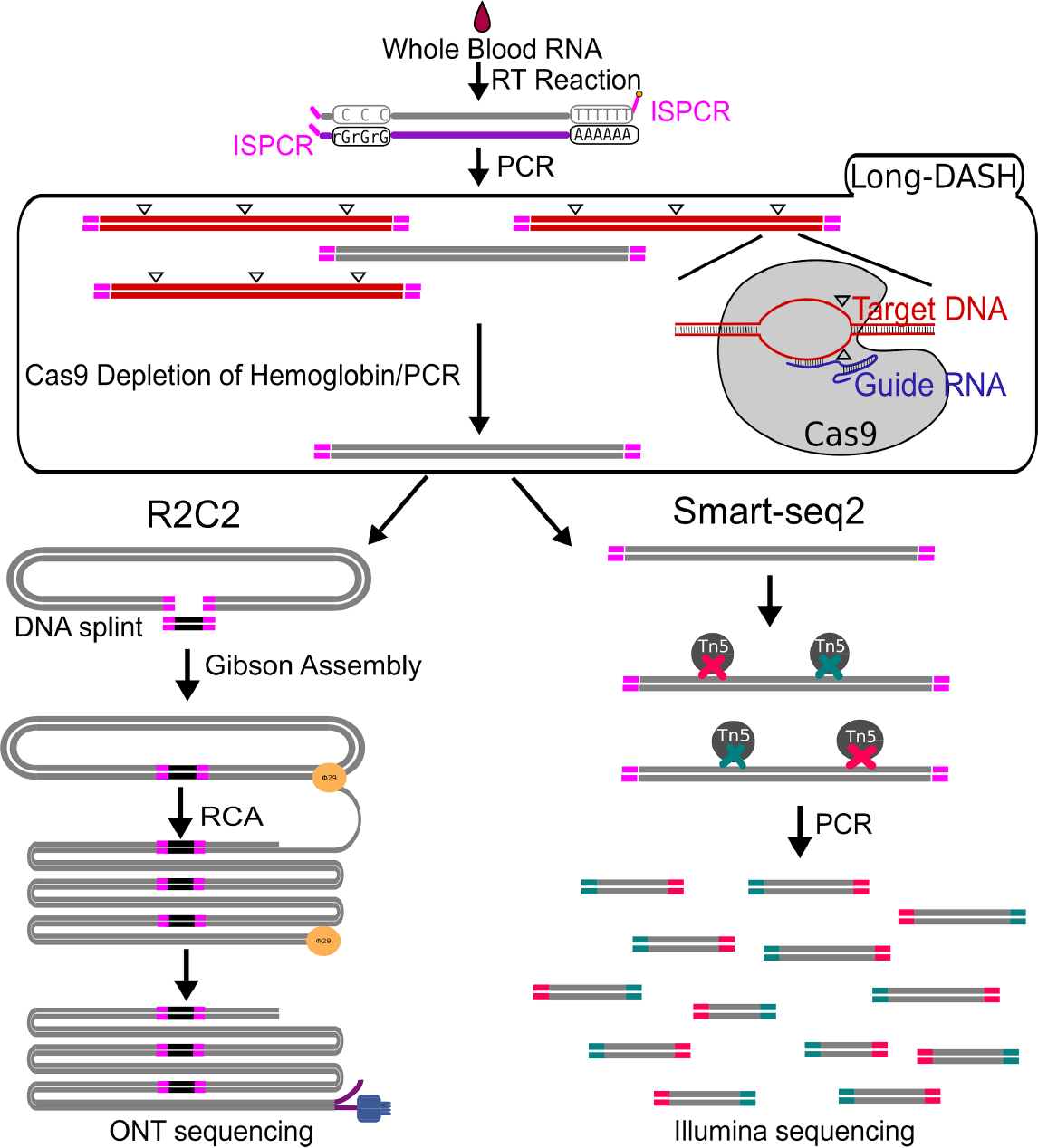
Schematic of Long-DASH. A) Whole Blood RNA is extracted and full-length cDNA is generated with the first half of the Smart-seq2 protocol. The cDNA is then depleted of hemoglobin transcripts using the recombinant S. pyogenes Cas9 protein bound to hemoglobin specific sgRNA which cuts hemoglobin cDNA molecules by introducing double strand breaks (△) in a sequence specific manner. The cut molecules can no longer be exponentially amplified with PCR, so a subsequent PCR step is performed to enrich for complete non-hemoglobin cDNA molecules. The resulting hemoglobin depleted cDNA pool is then sequenced using the ONT-based R2C2 library prep and the Illumina-based Smart-seq2 library prep.

**Figure 2:**
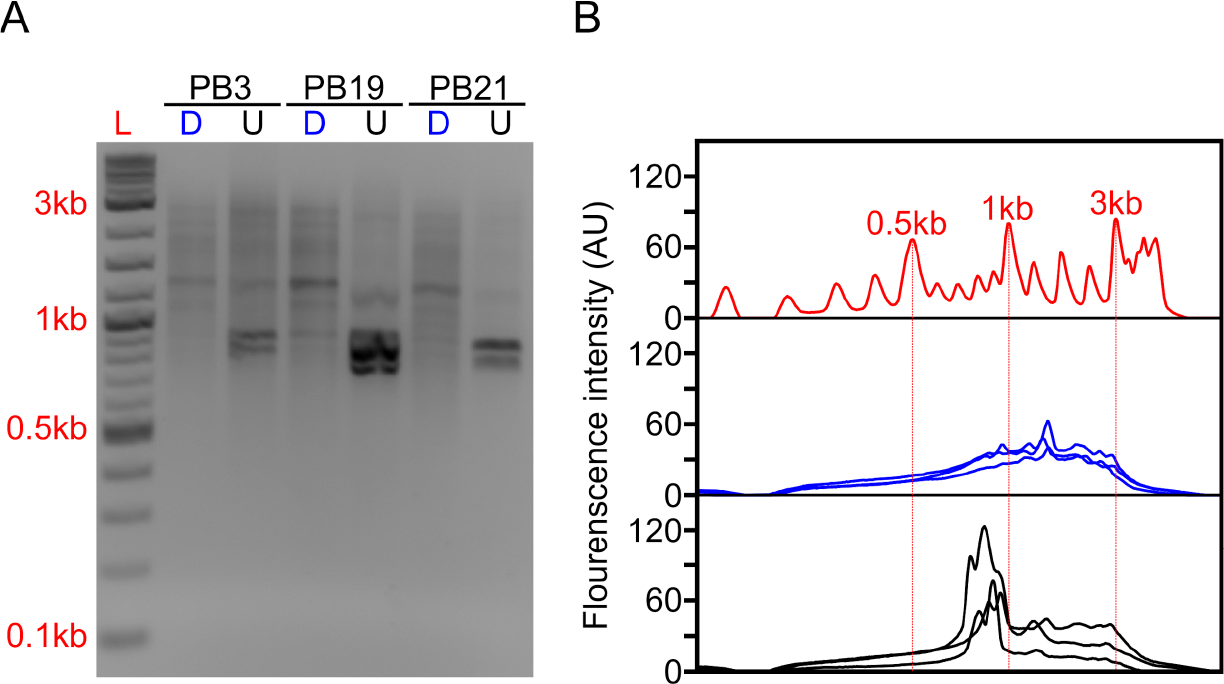
Long-DASH depletes hemoglobin from full-length cDNA. A) Depleted (D) or undepleted (U) cDNA was visualized on a 2% Agarose gel. DNA ladder (L) suggests highly abundant cDNA species - putatively hemoglobin around ~700bp. B) Fluorescence analysis of the gel by ImageJ (28) further emphasizes the difference between depleted (blue) and undepleted (black) cDNA pool. Select size markers in the DNA ladder (red) are indicated.

### Long-DASH is compatible with Smart-seq2 library preparation and does not distort cDNA composition

Next, we aimed to validate whether Long-DASH truly depletes hemoglobin transcripts in the cDNA pool and can be used for Illumina’s short-read RNA-seq sequencing platform. To show this we prepared independent Tn5 based Smart-seq2 sequencing libraries for each depleted and undepleted cDNA pool. We then sequenced the Smart-seq2 libraries on a multiplexed Illumina HiSeq X 2 × 151 bp run. We generated ~20 million reads for depleted and ~60 million reads for undepleted samples. By sequencing the undepleted samples deeper, we reasoned that the non-hemoglobin genes should receive equivalent read coverage in depleted and undepleted samples. This allowed us to perform a side by side comparison of the depleted and non-depleted samples to ensure no off-target effects.

First, we analyzed the resulting sequencing data using a custom kmer based approach to estimate the number of reads originating from hemoglobin transcripts. In the undepleted cDNA pools 48-68% of reads were scored as originating from hemoglobin transcripts. In depleted samples this was reduced to 1-4% reads (Fig. 3A).

**Figure 3:**
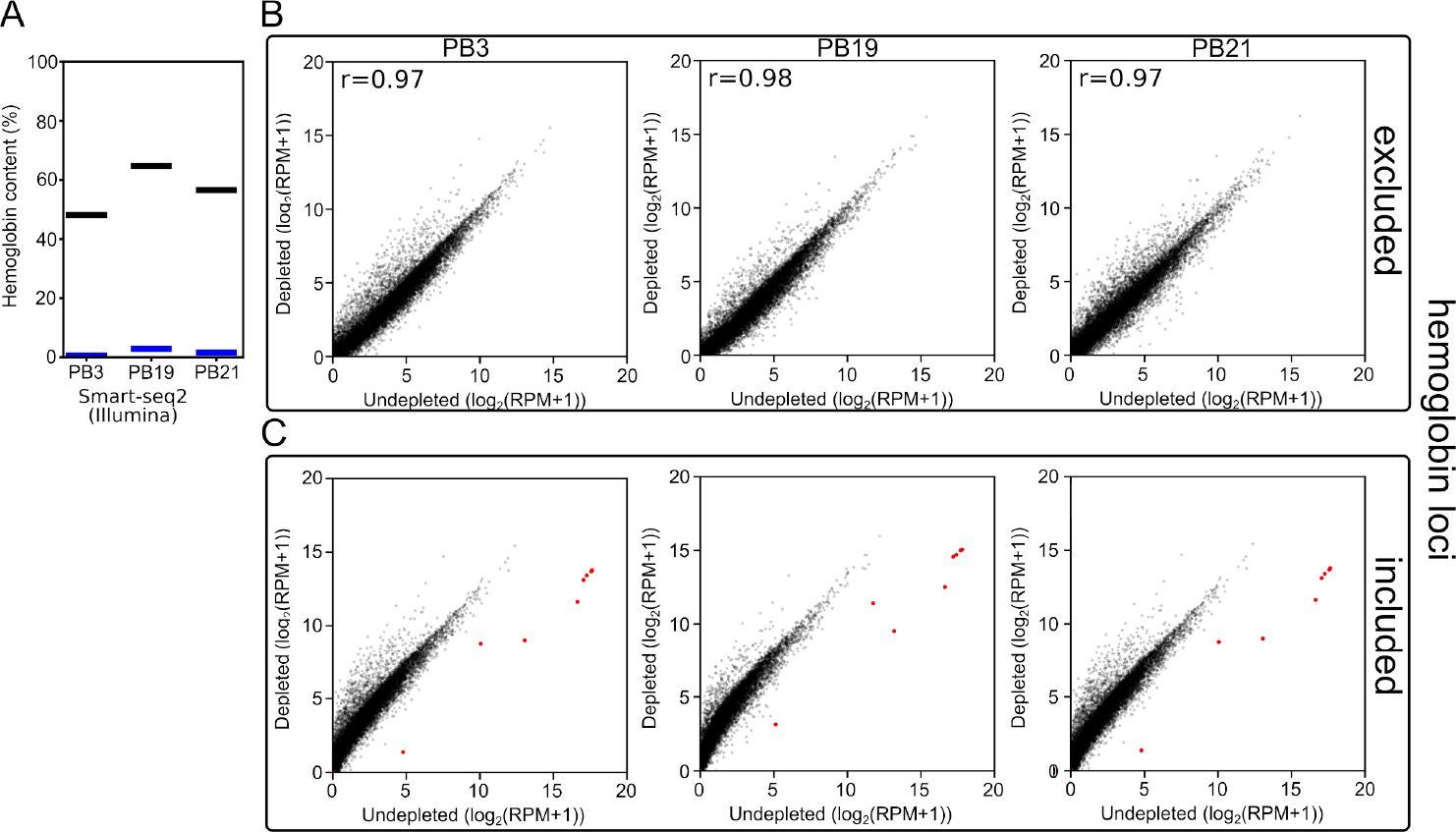
Long-DASH depletes hemoglobin from cDNA. A) Hemoglobin content was measured in Smart-seq2 (Illumina) libraries of depleted (blue) or undepleted (black) cDNA pools. B,C) Scatterplots comparing gene expression in undepleted and depleted Smart-seq2 libraries of PB3, PB19, and PB21 with reads aligning to hemoglobin loci (red) (29) either excluded (B) or included (C).

Second, to show that the depletion of hemoglobin transcripts did not distort the rest of the cDNA pool, we aligned the reads to the polar bear genome and quantified the expression of all previously annotated genes. Quantification of gene expression showed that the overall gene expression patterns were not dramatically distorted between depleted and undepleted samples. The three polar bear samples showed a Pearson r-value of 0.97-0.98 (Fig. 3B) when the gene expression values of depleted and undepleted samples were compared and reads aligning to hemoglobin loci were discarded. If reads aligning to hemoglobin loci are included in the analysis, the large number of reads aligning to the few hemoglobin loci in undepleted samples skews RPM calculations and shifts the overall correlation (Fig. 3C).

Overall, this showed that depletion of hemoglobin of full-length cDNA was successful, thereby freeing up the vast majority of sequencing reads to analyze the rest of the polar bear transcriptome.

### Long-DASH is compatible with full-length cDNA sequencing methods

Having established the compatibility of Long-DASH with the short-read RNA-seq assay, we investigated whether we could generate a long read data set from the depleted cDNA using our R2C2 approach. By incorporating R2C2 we can generate error-corrected full-length cDNA reads using long-read ONT sequencers. We used 5 partially multiplexed flowcells to generate ~3.8 million R2C2 consensus reads of 5 depleted cDNA pools – two Long-DASH replicates (R1 and R2) for PB3 and PB19 as well as a single Long-DASH run for PB21. The R2C2 reads we generated had a median accuracy of 94%, which is between 8-10% more accurate than standard ONT cDNA sequencing protocols (Table S1).

We also generated ~5,000 R2C2 consensus reads of undepleted cDNA from one polar bear which allowed us to compare hemoglobin content and consensus read length distributions between depleted and undepleted samples (Fig 4). In the undepleted sample, the majority of R2C2 reads were of two distinct lengths, both around 700 bp, likely representing the 79.3% of hemoglobin transcripts present in that sample. The 5 depleted samples showed a much more evenly distributed read length with a median hemoglobin content of 1.2% (0.6%-8.3%) (Fig. 4). Higher hemoglobin levels for R2C2 compared to Smart-seq2 based library preps (1-4%) using the same cDNA might be explained with R2C2 being somewhat biased towards transcripts between 500-1000 bp.

**Figure 4:**
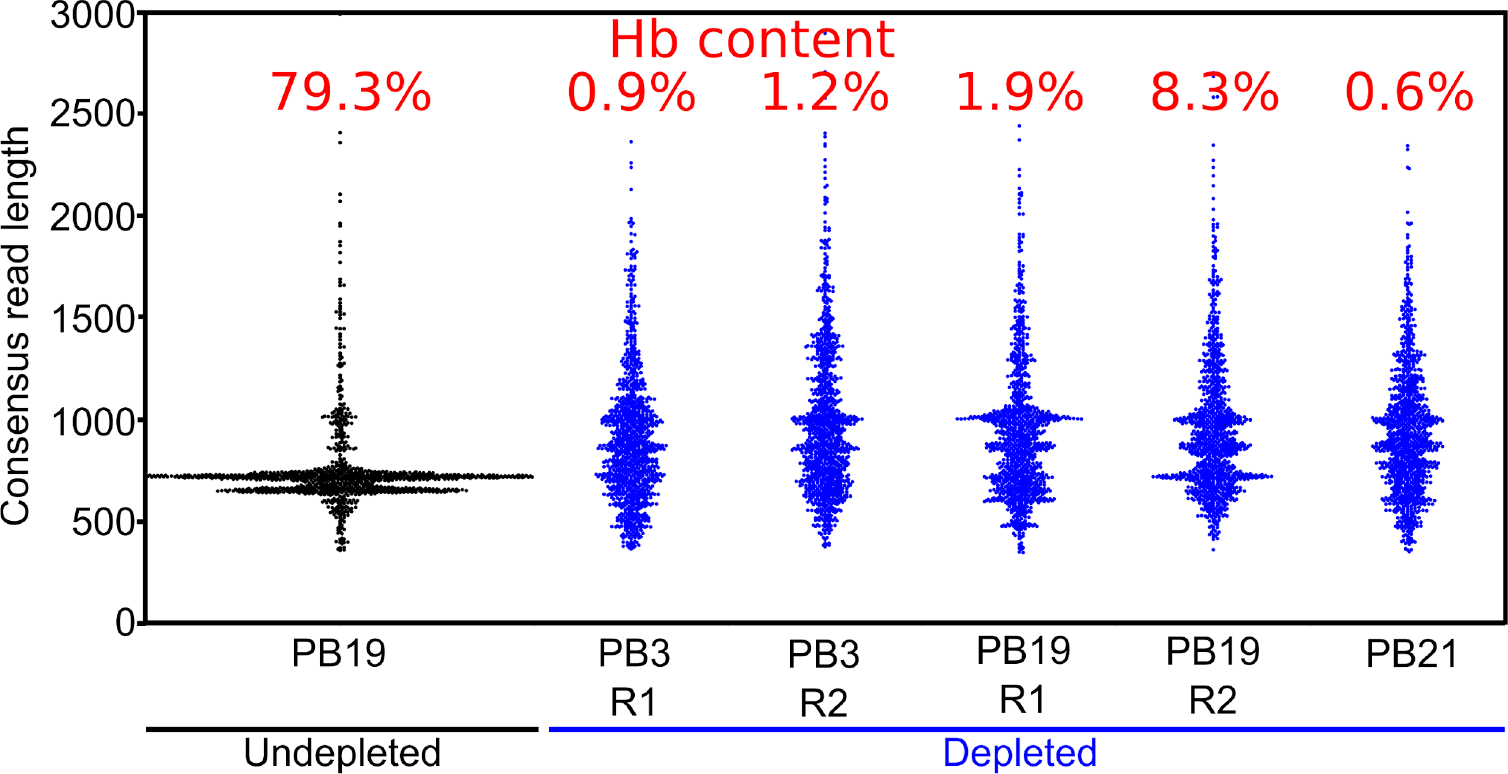
Long-DASH depletes hemoglobin from cDNA. Length distribution of R2C2 consensus reads is shown as swarmplots in the indicated samples. Independent Long-DASH replicates (R1 and R2) were performed for samples PB3 and PB19 but not PB21. Percent of Hemoglobin reads as determined with a kmer approach is given in red on top.

The median read length of the depleted samples was slightly below 1 kb which is in line with cDNA read length distributions published to date (30). This means that despite the less than ideal conditions for RNA integrity given difficult field conditions and the lag time between sample collection and processing, the analyzed RNA molecules were largely intact.

### R2C2 reads of depleted full-length cDNA can refine transcriptome annotations

Next, we generated high confidence isoform-level information from our full-length cDNA to refine the currently available polar bear transcriptome annotation. To this end, we analyzed our 3.8 million R2C2 consensus reads using the Mandalorion pipeline we previously developed (31). We aligned the R2C2 reads to the polar bear genome sequence (29) using minimap2. These alignments, together with previously known individual splice sites (29, 32), then serve as input into our Mandalorion pipeline which processes read alignments into high-confidence isoforms.

The Mandalorion pipeline first complements known splice sites with new splice sites it identifies de novo from R2C2 read alignments. It then groups R2C2 reads based on the splice sites they use. Pairs of transcription start sites (TSSs) and polyA sites are then determined for each group to identify full-length isoforms. Two additional processing steps were performed whereby isoforms were excluded if they were fully contained within longer isoforms or unspliced. This was to ensure removal of any non-full-length isoforms that may result from RNA degradation, as well as isoforms potentially caused by DNA contamination, respectively. In total, this analysis produced 5,831 high-confidence isoforms with a median accuracy of 99.1%.

We then classified these 5831 high-confidence spliced isoforms using the Squanti algorithm (33) that determines what relationship an experimentally determined isoform has to genes and isoforms in a reference annotation (Fig. 5). As a reference, we downloaded 28,880 known and predicted mRNA sequences from NCBI by selecting “RefSeq” and “mRNA” filters in the NCBI Nucleotide database most of which are based on the NCBI Ursus maritimus Annotation Release 100 catalog of polar bear mRNA sequences (34).

**Figure 5:**
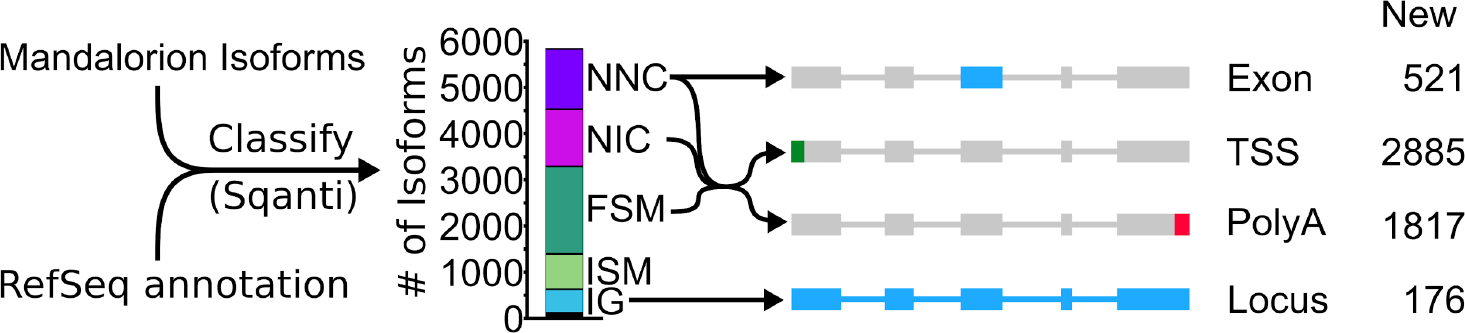
R2C2/Mandalorion Isoforms identify new features in the polar bear transcriptome. General workflow for comparing RefSeq mRNAs and Mandalorion isoforms is shown on the left. RefSeq mRNAs were aligned to the polar genomes using minimap2 and converted to gtf format to create a reference annotation. Isoforms determined by Mandalorion were then classified using this reference annotation using the sqanti_qc algorithm. Isoforms were classified as Novel_not_in_catalog (NNC), Novel_in_catalog (NIC), Full_splice_match (FSM), Incomplete_splice_match (ISM) and Intergenic (IG). New transcriptome features were then determined based on the minimap2 alignments of isoforms in the indicated categories.

1239 of the 5831 Mandalorion isoforms were classified as “novel_not_in_catalog” (NNC) which means that they overlapped a known gene but contained at least one unannotated splice site. In-depth analysis of this NNC group found that they contained a total of 521 new exons. 1301 isoforms were classified as “novel_in_catalog” (NIC), which means that they overlapped a known gene and used only annotated splice sites but at least once as part of a previously unannotated splice junction. In total we observed 2540 (1239 NNC and 1301 NIC) new isoforms with unannotated exon configurations. An additional 1893 isoforms were classified and “full_splice_match” (FSM) which means that their splice junctions matched an annotated isoform completely but it doesn’t mean that TSS and polyA sites also matched this isoform. In-depth analysis of the putative full-length NNC, NIC, and FSM isoform groups identified 2885 new TSSs and 1817 new polyA sites. Finally, 769 isoforms were classified as “incomplete_splice_match” (ISM) which means that they contain a subset of splice junctions of an annotated isoform. While these isoforms could represent real, shorter transcripts, they might also represent experimental artifacts so we excluded them from TSS and polyA analysis.

Considering RefSeq mRNA sets are in part based on deep short-read data and computational annotation, we did not expect to discover many entirely new gene loci. However, 509 of the 5831 isoforms were classified as “intergenic” (IG) which means that they did not overlap with any annotated gene locus. By determining which of these isoforms overlapped with each other, we identified 176 new gene loci.

Overall, this analysis dramatically refined our isoform-level knowledge of the whole blood polar bear transcriptome (Fig. 5). To make this knowledge straightforward to use for future analysis, we have generated a gtf annotation file containing RefSeq mRNA entries merged with our R2C2/Mandalorion isoforms.

How these new isoforms and isoform features have improved the current annotation can be seen clearly in these three following examples. In the RBX1 gene, we discovered 10 new isoforms containing several new TSSs and polyA sites, several were associated with new terminal first or last exons (Fig. 6A). In the GMFG gene, we similarly identified new isoforms containing unannotated internal and terminal exons, intron retention events, TSSs, and polyA sites (Fig. 6B). Finally, we discovered a new gene locus that contains two isoforms and is entirely absent in the polar bear RefSeq mRNA set. However, aligning the the two isoforms to the Panda genome (35) resulted in unique matches to the CCDC72 gene (Fig. 6C).

**Figure 6:**
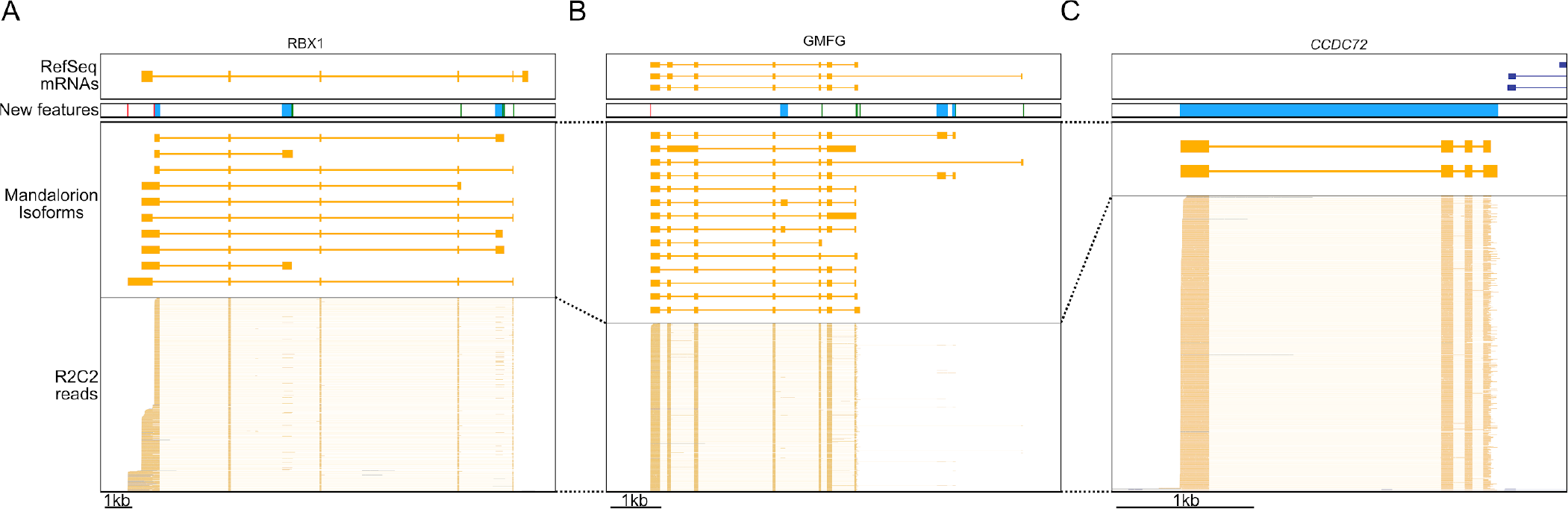
R2C2/Mandalorion refine transcriptome annotations. Genome Browser views of the RBX1 locus (A), the GMFG locus (B), a locus likely corresponding to the CCDC72 gene not yet included in the RefSeq mRNA set (C). From top to bottom, 1) RefSeq mRNAs alignments, 2) new features based on Mandalorion isoforms (green: TSS, red: polyA site, blue: new exon or locus), 3) Mandalotion isoforms, and 4) R2C2 reads. Plus strand alignment are in blue, minus strand alignments in orange.

## Discussion

To better understand how humans and environmental perturbations impact threatened or endangered species, it is critical to understand changes in transcriptome dynamics. Fluctuations at the molecular and cellular level are sensitive indicators of environmental change (36, 37); they are analogous to veterinary medicine where blood transcriptomes serve as proxies for identifying health status, disease, and exposures to environmental toxicants (38–41). Changes at the transcriptome level may also be useful indicators of ecological specialization, and therefore useful to design strategies for species management and conservation (42). However, existing approaches to generate transcriptome data from whole blood RNA are either specifically designed for short-read sequencing (DASH) or human samples (commercial hemoglobin depletion kit like GLOBINclear) and therefore lack a cost-effective approach for analyzing isoform-level transcriptomes of non-model organisms.

Any study investigating whole blood transcriptomes using short- or long-read sequencing will greatly benefit from the Long-DASH method. Long-DASH effectively and economically depletes hemoglobin transcripts from whole blood full-length cDNA which can then be sequenced with short- or long-read sequencing. We validated Long-DASH by depleting hemoglobin transcripts from polar bear whole blood cDNA pools and generated deep short-read Smart-seq2 RNA-seq data as well as 3.8 million R2C2 full-length cDNA consensus reads. We processed the 3.8 million full-length R2C2 reads to identify close to 6000 high confidence isoforms which we then used to refine and improve the polar bear whole blood transcriptome annotation.

In addition to polar bear hemoglobin transcripts, the sgRNAs designed for this study will also target human hemoglobin transcripts making them useful for basic research as well as clinical applications in cancer biology and disease diagnosis (Fig. S1) (43–46). Further, the sgRNA sequences used in Long-DASH can be easily adapted to any organisms or gene. The ease and adaptability places Long-DASH at an advantage over “as-is” commercial kits like GLOBINclear (Ambion), which promises >95% of depletion of human and mouse hemoglobin transcripts, but fails to efficiently deplete hemoglobin transcripts from pig whole blood RNA samples (22).

Since cDNA can be generated from femtogram levels of polyA-RNA, Long-DASH requires very little RNA input compared to RNA pull-down methods. This allows the investigator to gather small samples, or only process small aliquots of existing samples, thereby maximizing the usefulness of each sample collection and minimizing harm to animals.

Going forward, the Long-DASH depletion method and the R2C2 long-read sequencing method will form a very powerful combination for transcriptome analysis and annotation from whole blood samples and beyond. The transcriptomes of many tissues contain several highly abundant transcripts that represent >50% of all transcript molecules (47). A set of sgRNAs targeting any abundant transcripts can be easily generated, making Long-DASH conducive for surveying other tissues as well. Specifically, depleted full-length cDNA libraries can be sequenced using our R2C2 method, which currently represents the most powerful combination of throughput and accuracy in the long-read sequencing field. Our most recent R2C2 run emphasizes this by generating ~1,000,000 R2C2 reads at a median accuracy of 97.5% on a single ONT MinION flowcell at a cost of ~$650 (Table S1). This represents an increase in accuracy of >10% over standard ONT cDNA sequencing and 10-times more complete reads than the PacBio Sequel at the same cost. Combining our Long-DASH and R2C2 methods therefore brings the exhaustive annotation of non-model organisms within reach.

## Materials and Methods

### Sample Collection/RNA Extraction from Whole Blood

Permits for field operations and animal care were provided by the Government of Greenland (Permit numbers 2015-110281 and 2017-5446). Polar bear whole blood samples were collected in PAXgene Blood RNA tubes (PreAnalytiX GmbH, BD Biosciences, Mississauga, ON, Canada). Total RNA was isolated from whole blood (2.5mL) thawed at room temperature for 2 hours prior to using the PAXGene RNA extraction kit (Qiagen, Chatsworth, CA, USA) according to manufacturer’s protocol. All samples were DNAse (Qiagen) treated and eluted in 50μL. The RNA yield and purity were accessed using a NanoDrop 8000 UV Spectrophotometer (Thermo Fisher Scientific, Waltham, MA, USA). RNA quantities ranged from 110 - 310 ng/μL and the A260/280 ratio values were > 2.0. Human whole blood RNA was purchased from Zyagen Labs (NC1453913).

### Full-length cDNA Generation

RNA was reverse transcribed (RT) using Smartscribe Reverse Transcriptase (Clonetech). We generated full-length cDNA using a modified Smart-seq2 approach (27). During the RT reaction a template-switch oligo and an oligodT primer was used to select for polyA+ RNA (Table S2). The RT reaction was performed in 10 μL reactions with an input of 70 ng of RNA and took place at 42°C for 1 hour. After cDNA synthesis, 1 μL of 1:10 dilutions of RNAse A (Thermofisher) and Lambda Exonuclease (NEB) were added and incubated at 37°C for 30 minutes. Following the incubation, an amplification step was performed in 25 μL final volumes using KAPA Hifi ReadyMix 2X (KapaBiosystems) containing 1 μL of the ISPCR primer (10 μM) primer. Samples were incubated at 95°C for 3 minutes, followed by 12 cycles of (98°C for 20 s, 67°C for 15 s, and 72°C for 4 minutes), with a final extension of 72°C for 5 minutes. Samples were purified using Agencourt AMPure XP SPRI beads (Beckman Coulter) and eluted at 25 mL. The final cDNA product was then visualized on an agarose gel to confirm distribution (Fig. 2).

### In-vitro Preparation of CRISPR/Cas9

SpCas9-2xNLS was purified based on the protocol described in (48). Briefly, a plasmid encoding His6MBP-SpCas9-2xNLS (Addgene plasmid #69090) was transformed into Rosetta2(DE3) *E. coli* cells. Cultures were grown at 37ºC in 2YT medium with shaking until they reached an OD_600_ of ~0.6, and then placed on ice for 5 minutes before adding IPTG to a final concentration of 0.25 mM; cultures were then grown overnight at 18ºC with shaking. Cell pellets were harvested by centrifugation, and then lysed in an Avestin cell extruder in Ni-A buffer (20 mM Tris pH 8.0, 500 mM NaCl, 5% vol/vol glycerol, 25 mM imidazole) with EDTA-free protease inhibitors (Pierce). Clarified supernatants were purified by gravity column on Ni-NTA agarose (QIAGEN) using Ni-A buffer to load and wash, and Ni-B buffer (20 mM Tris pH 8.0, 500 mM NaCl, 5% vol/vol glycerol, 250 mM imidazole) to elute. Peak fractions were concentrated in an Amicon Ultra spin concentrator with a 30 kDa molecular weight cutoff at 4ºC, and then loaded onto a 50 mL HiPrep Desalting Column (GE Healthcare) pre-equilibrated with 17% IEX-B (IEX-A buffer: 20 mM HEPES pH 7.5, 150 mM KCl, 5% vol/vol glycerol; IEX-B 20 mM HEPES pH 7.5, 1 M KCl, 5% vol/vol glycerol). The flow-through was then loaded onto a 2 mL HiTrap SP column (GE Healthcare) in 17% IEX-B buffer. After thoroughly washing the column in 17% IEX-B, the protein was eluted with a linear gradient from 17-50% IEX-B. Peak fractions were pooled and loaded onto a Superdex 200 16/60 column (GE Healthcare) pre-equilibrated in 20 mM HEPES pH 7.5, 150 mM KCl, 1 mM DTT, 10% vol/vol glycerol. Peak fractions were concentrated in an Amicon Ultra spin concentrator with a 30 kDa molecular weight cutoff at 4ºC until a concentration of 40 ſM, which was estimated using the calculated molar extinction coefficient of 120,575 M^−1^ cm^−1^. The protein was aliquoted into small volumes (10 ſL), quick frozen in liquid nitrogen, and stored at −80ºC.

### sgRNA Design and Construction

Other studies have shown that sgRNAs designed between 17-20 bp showed increased efficacy (49, 50). As a result, the sgRNAs were designed between 17-20 bp in length. sgRNAs were designed to target hemoglobin transcripts in human and polar bear. A multi-sequence alignment was performed on the human and polar bear annotated HBA and HBB gene transcripts to find conserved regions using the Clustal Omega tool (51–53) (Fig. S2). Regions with high homology were chosen for sgRNA design. sgRNAs that did not share complete homology were designed to contain degenerate bases to ensure compatibility across species using the same sgRNA (Fig. S3). sgRNA specificity was determined by using BLAST (54). One sgRNA was designed even though the N-GG (PAM motif) had been lost in the human but was still kept in the pool for the polar bear depletion (Fig. S3). A total of 16 sgRNAs were designed to target alpha and beta hemoglobin transcripts. The target oligos were then constructed into sgRNAs as previously described (49). Single stranded oligos were designed to contain a T7 promoter attached to each sgRNA sequence (IDT) followed by the first 22 bases of the tracrRNA sequence (Fig. S3). The complementary tracrRNA and single stranded oligo were annealed and extended to form a dsDNA product containing the T7-sgRNA and tracrRNA template. The template was then used for in-vitro transcription using the HiScribe T7 High Yield RNA synthesis kit (NEB). The in-vitro transcription reaction was carried out at 37°C for 16 hrs. The in-vitro transcribed RNA was then purified using MEGAclear Transcription Clean-Up Kit (Invitrogen). The final sgRNA product was then checked for purity and quantified using NanoDrop 8000 UV Spectrophotometer (Thermofisher). All sgRNAs were then pooled at equal molar concentrations and stored in single-use aliquots at −80°C.

### CRISPR/Cas9 Treatment

Since it has been predicted that human whole blood samples can contain up to ~50-80% of hemoglobin transcripts of the total sample (20, 21), we calculated the ratio of sgRNA and Cas9 molar amounts to sample based upon this assumption. According to the DASH protocol it was determined that 150-fold of Cas9 and 1500-fold of sgRNA should be sufficient (24). All cDNA samples were quantified by Qubit using the dsDNA HS assay kit (Thermofisher) to calculate the molar amounts. To calculate the expected molar amounts we use the following formula:

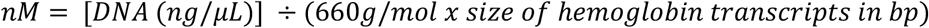

Once the molar amounts were determined, the ribonucleoprotein (RNP) complex was formed by adding the 150-fold Cas9 and 1500-fold sgRNA excess amount with 1.0 μL of the 10X Cas9 Buffer (final concentration 50 mM Tris pH 8.0, 100 mM NaCl, 10 mM MgCl_2_, and 1 mM TCEP) and incubated for 25°C for 10 minutes. Following the 25°C incubation, the calculated cDNA amount was then added (final volume of 10 μL) and incubated at 37°C for 4 hrs to overnight. After the Cas9 depletion, 1 μL of Proteinase K and RNAse A were added to inactivate the Cas9 and remove excess sgRNAs from the reaction and incubated at 37°C for 15 minutes and 95°C for 15 minutes. It is critical that the Proteinase K is deactivated properly as the samples are immediately used for amplification. Treated samples were PCR amplified (95°C for 3 minutes, followed by 13 cycles of (98°C for 20 s, 67°C for 15 s, and 72°C for 4 minutes) followed by a final extension of 72°C for 5 minutes). PCR was performed using KAPA Hifi ReadyMix 2X (KapaBiosystems) and 1 μL of the (10 μM) ISPCR primer. The amplified product was then purified using SPRI beads to remove everything below 500 bp. Selecting against cDNA below 500 bp ensured that all cut hemoglobin products were removed before making the Tn5 libraries. The depleted cDNA product was visualized on a 1-2% agarose gel to confirm depletion. Once confirmed, the depleted cDNA product was then prepped for either Illumina or Nanopore sequencing.

### R2C2 Library Preparation and ONT sequencing

To prepare R2C2 libraries ~30 ng of the depleted cDNA was used. The R2C2 libraries were made as previously described (25) with modifications. DNA splint was generated with a primer extension reaction containing 1ul each of 100uM of UMI_Splint_Forward and UMI_Splint_Reverse primers (Table S2) and KAPA Hifi ReadyMix 2X (KapaBiosystems) incubated at (95°C for 3 min, 98°C for 1 min, 62°C for 1 min, 72°C for 6min) and purified using SPRI beads. An equal concentration of splint to cDNA were combined (30ng of depleted cDNA and 30ng of our (~200 bp) DNA splint). The full-length cDNA was then circularized using the 2X NEBuilder Hifi DNA Assembly Mix (NEB). The reaction took place at 50°C for 1 hour per manufacturer protocol. Once the full-length cDNA was circularized, linear ssDNA and dsDNA was digested by adding 3μL each of Lambda Exonuclease, Exonuclease I and Exonuclease III (all NEB) and incubated at 37°C overnight. We performed the longer incubation overnight to ensure complete digestion. After digestion, the sample was further purified using SPRI beads and eluted in 30μL of ultrapure water. 30μL of sample was then split into three reactions containing 10μL each for the Phi29 amplification. The Phi29 amplification took place in a reaction volume of 50μL containing 5μL of 10X Buffer, 2.5μL of 10uM each dNTPs, 2.5μL of random hexamers (10uM), 29μL of ultrapure water and 1μL of Phi29 Polymerase. The Phi29 reactions were incubated at 30°C for 16 hrs, 65°C for 15 minutes and held at 4°C. All three samples were pooled together and ultrapure water was added to make up the final volume to 300μL. The product was purified using SPRI beads with a 1:0.5 sample to bead ratio. This ratio was chosen as it removed all fragments < 2000 kb. The sample was then eluted in 90μL of ultrapure H20, 10μL of NEB2 Buffer (NEB) and 3μL of T7 endonuclease (NEB) and incubated at 37°C for 2 hrs to ensure complete debranching of the Phi29 product. The eluted sample was again purified using SPRI beads with a 1:0.5 sample to bead ratio. The product was eluted in 30μL and quantified using Qubit dsDNA HS kit (Thermofisher). The length distribution was verified on a 1% agarose gel prior to performing the ONT library prep.

For the library preparation ~1-2 μg of the final R2C2 product was converted into a ONT compatible library using the SQK-LSK109 kit according to ONT instructions with minor modifications. First during the End Repair and A-tailing reaction we performed incubations for 30 min each at 20°C and 65°C instead of 5 min each. Second we adjusted the ligation reaction time to 30 minutes at room temperature instead of 10 minutes per the protocol. We also found that loading between ~200-500 ng of the final library onto the flowcell was the most optimal. Loading more library resulted in severe loss in throughput as can be seen for the R2C2 runs PB3_depleted_R1 and PB19_depleted_R1 (Table S2). R2C2 libraries were sequenced on a MinION device using the 48hr sequencing protocol using the FLO-Min106 R9.4 Rev D chemistry flowcells. All reads were basecalled with Albacore v2.1.3.

### Smart-seq2 Library Preparation and Illumina Sequencing

Illumina libraries of the depleted and non-depleted samples were prepared using a tagmentation based method using our own Tn5 (26). The Tn5 enzyme was custom loaded with Tn5ME-A/R and Tn5ME-B/R adapters (Table S2). The Tn5 reaction contained 5μL of the full-length cDNA product, 1μL of the loaded Tn5 enzyme, 10μL of ultrapure water and 4μL of the 5X TAPS-PEG buffer and incubated at 55°C for 7 minutes. After incubation, 5μL of 0.2% of Sodium Dodecyl Sulfate (SDS) was added to the product to inactivate the Tn5 enzyme. Due to the Tn5 generating gaps, 5μL of the Tn5 product had to be nick translated at 72°C for 5 mins. The Tn5 product was then amplified using KAPA Hifi Polymerase (KAPA) with 10 cycles of PCR using (98°C for 10 s, 63°C for 30 s, 72°C for 2 min) with a final extension at 72°C for 5 min. The final reaction volume was 25μL and contained 0.5μL KAPA Hifi Polymerase (KAPA), 5μL of 5X Buffer, 0.8μL of dNTPs (10mM each), 11.7μL of ultrapure water, 5μL of the nick-translated product and 1μL each of Nextera_Primer_A and Nextera_Primer_B primers (Table S2). The amplified Tn5 libraries were then size selected from 300 - 800 bp on a 2% EX E-gel (Thermofisher) and purified using QIAquick gel extraction kit (Qiagen). The libraries were then pooled at equal concentrations and ran on a HiSeq × 2×151 bp run.

### R2C2 read processing and isoform analysis

R2C2 consensus reads were generated from raw reads using the C3POa pipeline (https://github.com/rvolden/C3POa). C3POa identifies subreads in the raw reads and uses poaV2 (55) and racon (56) to determine a more accurate consensus of these subreads. The consensus reads were then aligned to the polar bear genome (29) using minimap2 (57) using standard setting and the *‘-ax splice’* flag. The resulting sam files are converted to psl files using samtools (58) and jvarkit samtopsl utility (59).

The resulting psl, sam, and fasta files of all depleted samples were merged and used as input into the Mandalorion (https://github.com/rvolden/Mandalorion-Episode-II) pipeline to determine isoforms. To accomodate issues regarding RNA degradation and genomic DNA contamination, we integrated two new optional filter into Mandalorion. We implemented the filtering of isoforms that are entirely contained within one other isoform, which indicates degraded input RNA molecules, and the filtering of unspliced isoforms which might stem from DNA contamination.

Accuracy of R2C2 reads and Mandalorion isoforms were determined using alignments in sam format containing md-strings and a custom script that calculates

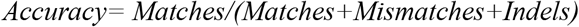

### Smart-seq2 read processing

Paired fastq files were downloaded from basespace and aligned to the polar bear genome using STAR with standard settings. The STAR index for the polar bear genome was built without a transcriptome reference because the gff file provided by (29) did not conform to gff standard (no “exon” features) and could therefore not be used. Read alignments in ordered bam format were converted to psl as described above.

### Hemoglobin and gene expression quantification

Hemoglobin content was determined through a kmer based counting method using a custom script. In short, all possible 10nt kmers were extracted from the sequence of hemoglobin alpha and beta transcripts. The presence of these kmers were then determined in each R2C2 or Smart-seq2 read from depleted and undepleted samples. Cutoffs for read assignments to hemoglobin were then determined by also analyzing R2C2 and Illumina reads of the GM12878 cell line which does not express hemoglobin.

Gene expression was determined using Smart-seq2 (Illumina) read alignments in psl format and a custom script. Reads aligning to hemoglobin loci were not counted towards total aligned reads in the RPM calculations.

Both script are available are available at https://github.com/christopher-vollmers/PB_scripts

## Data Visualization

Schematics were prepared using inkscape (https://inkscape.org). All others figures were prepared using python/matplotlib/numpy/scipy (60–63)

## Data Availability

All Illumina and ONT raw read data is available at SRA under Bioproject accession PRJNA514749.

## Acknowledgements

We thank Professor Rebecca Dubois and her lab at UC Santa Cruz for producing and providing us with the recombinant Tn5 enzyme used in this study and Professor Carrie Partch and her lab at UC Santa Cruz for producing and providing us with the recombinant Cas9 enzyme used in this study.

We acknowledge funding by the National Human Genome Research Institute/National Institute of Health Training Grant 1T32HG008345-01 (to A.B. and R.V.), the 2017 Hellman Fellowship (to C.V.), National Science Foundation grant DEB 1754451 (to. B.S.), funding for the field work provided by Environmental Protection Agency (Ministry of Environment and Food of Denmark) DANCEA Program and the Greenland Institute of Natural Resources.

## Author Contributions Statement

AB developed the Long-DASH method. AB, MS, KL, BS, and CV conceived and designed research. KL collected polar bear samples. AB and MS processed samples. AB, RV, and CV analyzed data. AB and CV wrote manuscript draft. AB, MS, RV, KL, BS, CV edited the manuscript.

## Supplementary Data

**Figure S1:**
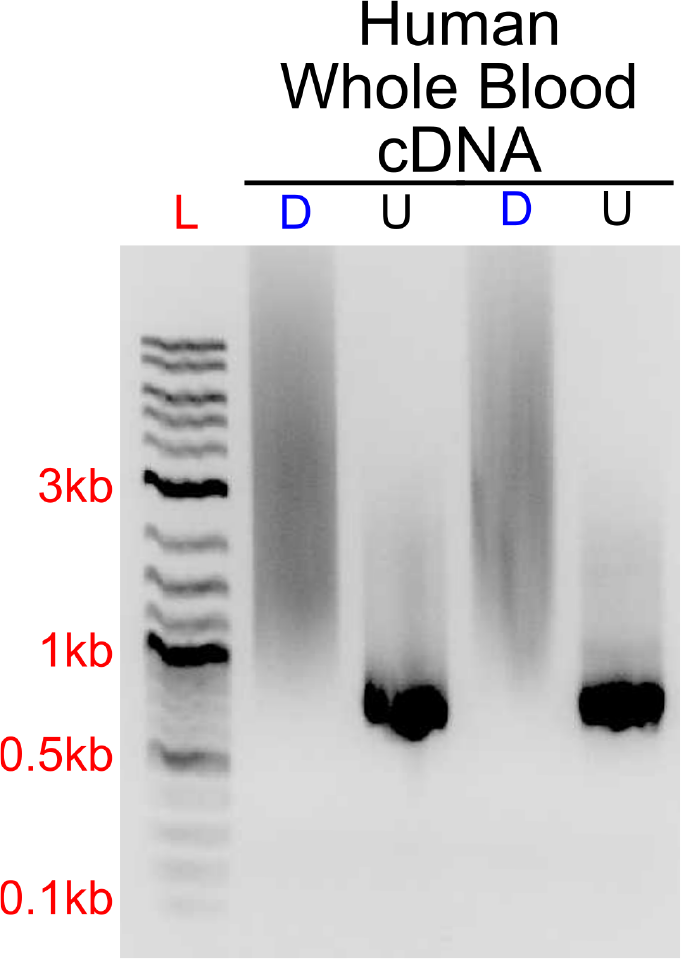
Long-DASH sgRNA also depletes human hemoglobin transcripts from full-lengthcDNA. Technical replicates of depleted (D) or undepleted (U) human whole blood cDNA were visualized on an agarose gel. DNA ladder (L) suggests highly abundant cDNA species - putatively hemoglobin around ~700bp.

**Figure S2.**
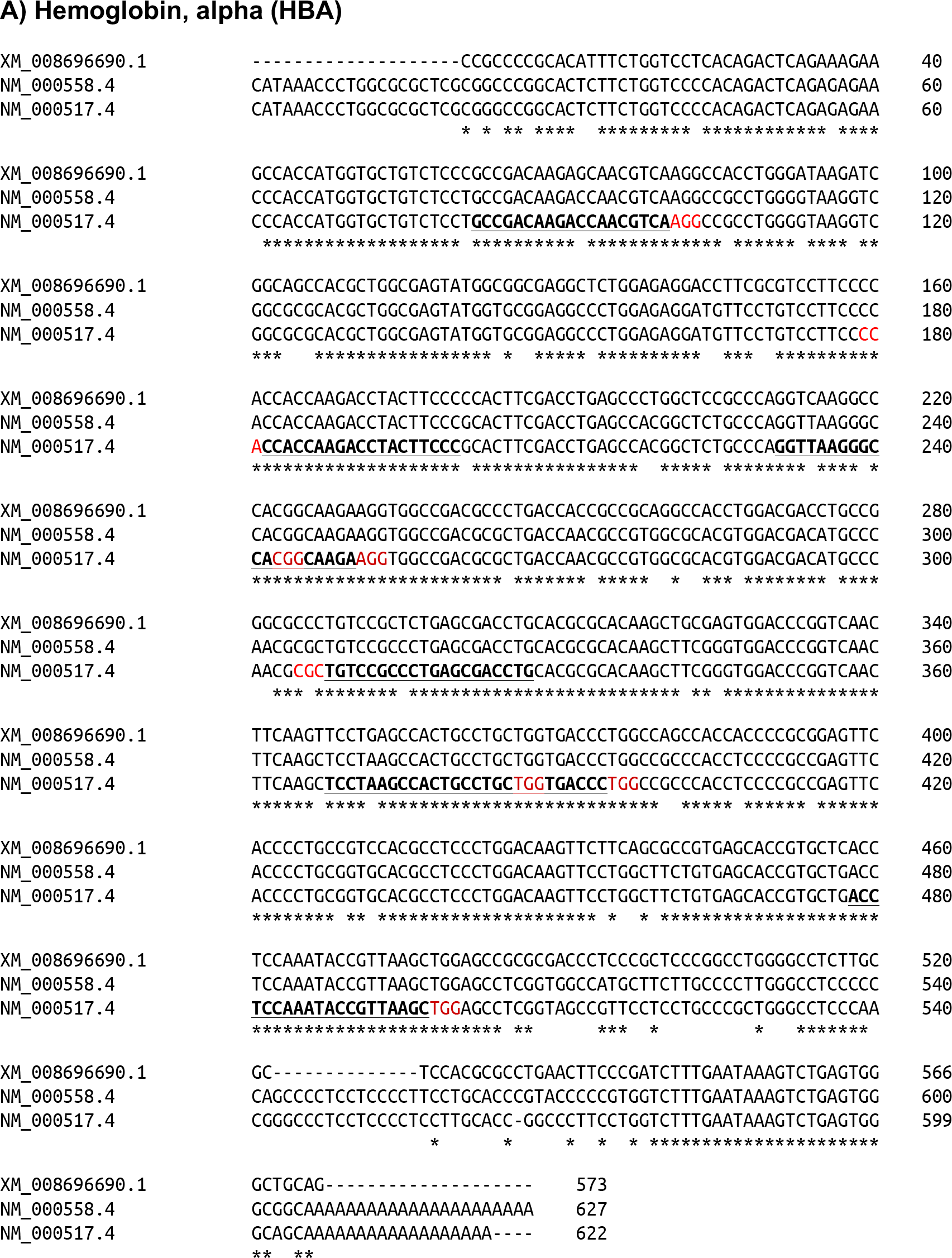
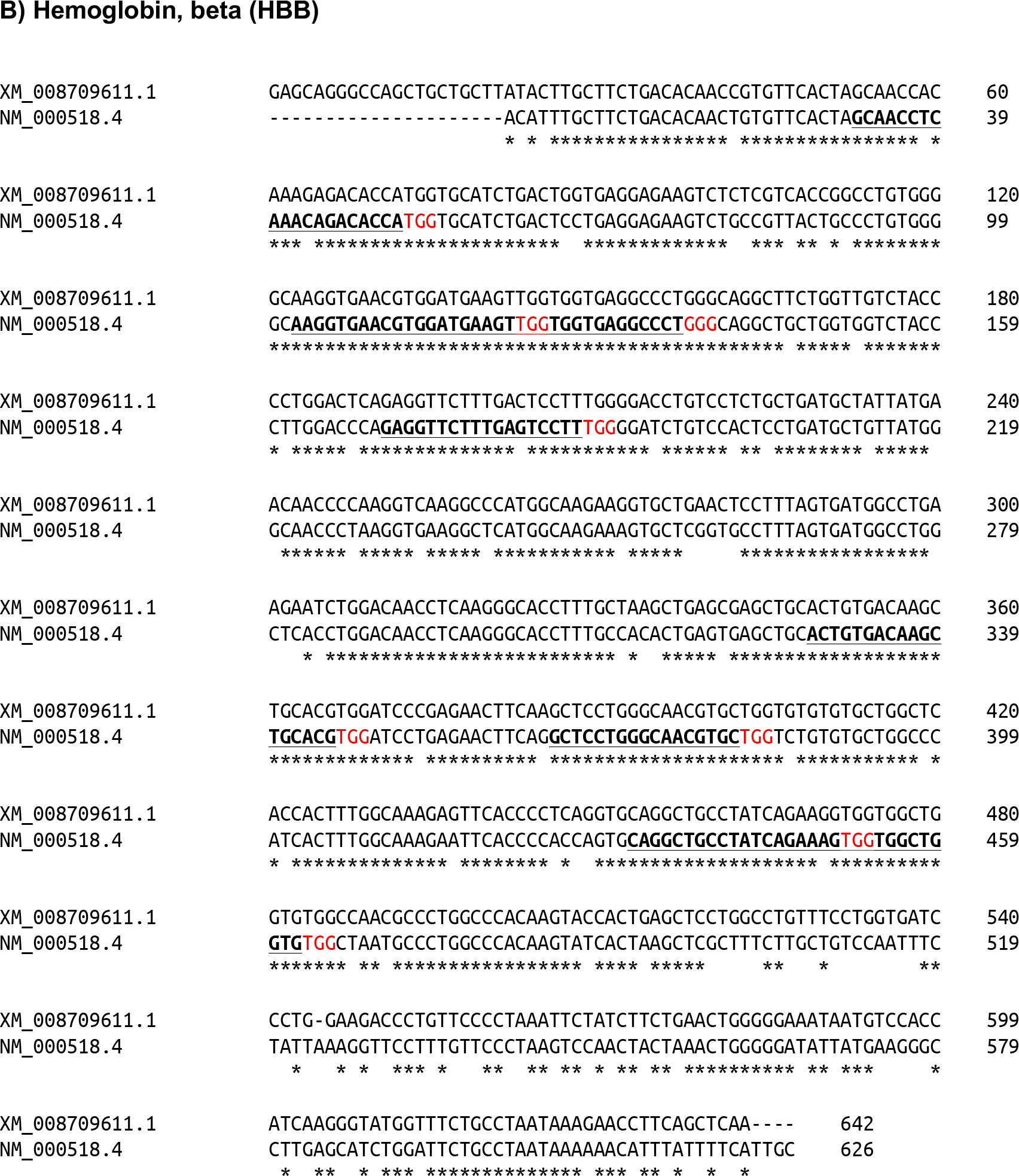
Alignment of orthologous HBA and HBB mRNA sequences in human and polar bear. Multi-sequence alignment from Clustal Omega v1.2.4. * indicates a match. Underline and bold indicates target sequences used for sgRNA design for Globin depletion. Red indicates (N-GG) PAM sequence a) Hemoglobin, alpha (HBA) b) Hemoglobin, beta (HBB).

**Figure S3.**
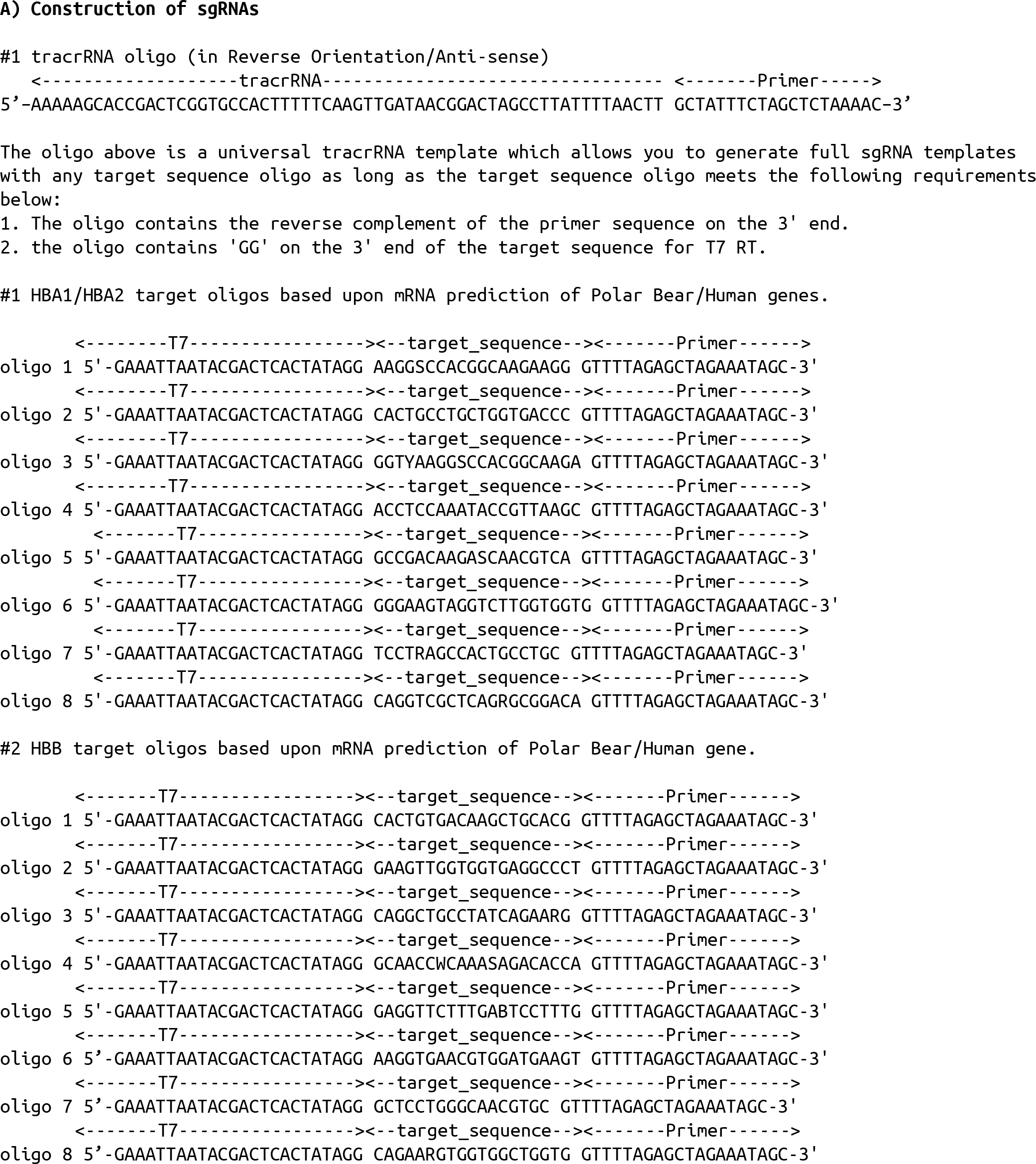
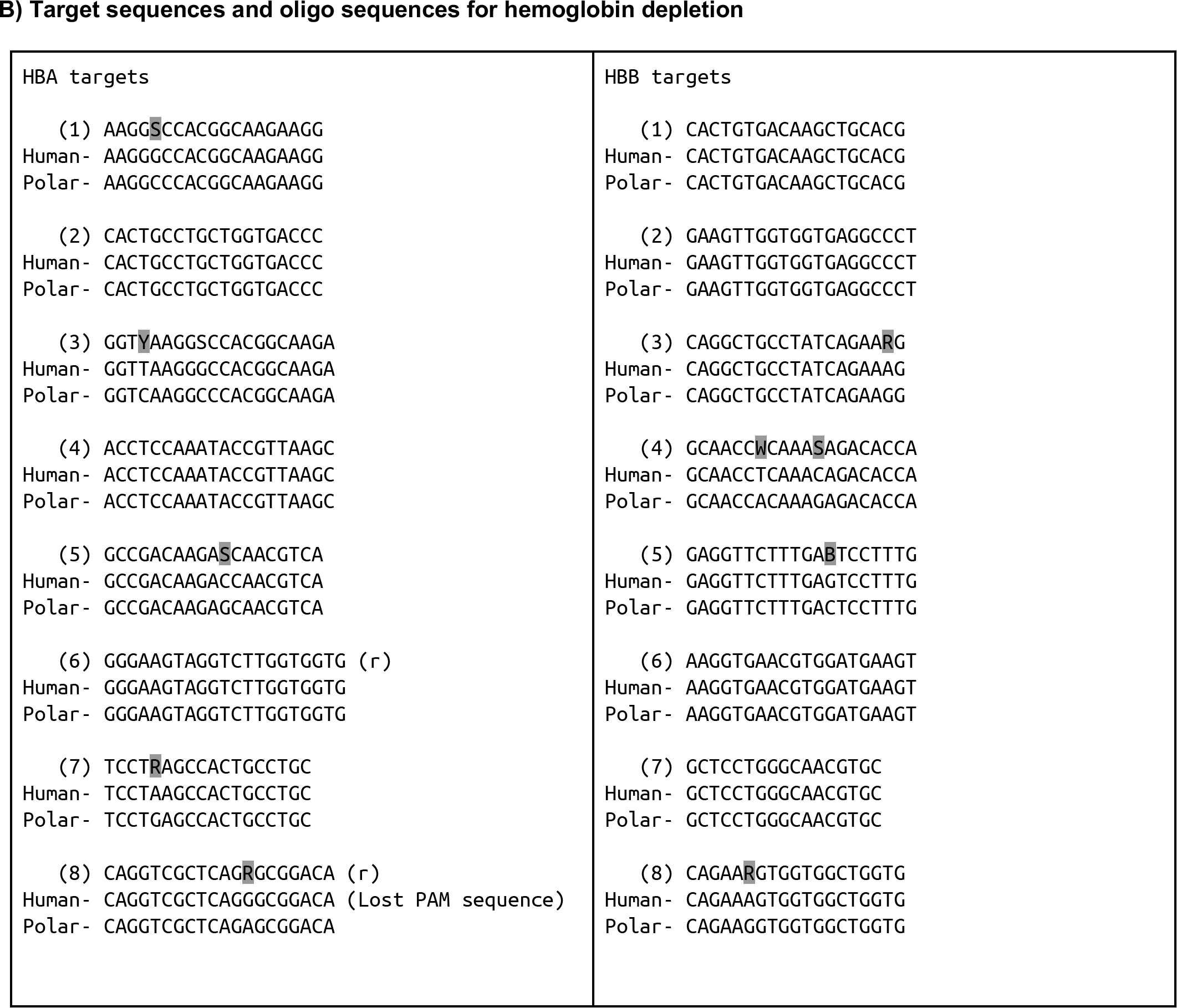
sgRNA design and construction. Oligonucleotides designed for hemoglobin depletion from full-length cDNA. Oligos were chosen to deplete Hemoglobin mRNA transcripts from Human and Polar Bear whole blood. A) To generate sgRNAs a template free PCR was performed to anneal the tracrRNA oligo to an oligo containing the target sequence to generate a full-length oligo. The full-length oligos were converted into sgRNA templates using in-vitro transcription. B) Target oligos used for generating sgRNAs. Degenerate bases are highlighted in grey. (r) indicates reverse orientation

**Table S1:** High-throughput sequencing runs and characteristics. For R2C2/ONT MinION runs, fully processed R2C2 read numbers and median accuracies are given. Some R2C2/ONT MinION runs were multiplexed, sometimes with samples unrelated to this study. Samples PB19_depleted_R2 and PB3_depleted_R2 represent the current output of the R2C2/ONT MinION combination

| Sample Name | Sequencer | Run Type | Library Prep | Read Number | Median Accuracy |
| --- | --- | --- | --- | --- | --- |
| PB3_depleted | Illumina HiSeqX | 2X151 | Smart-seq2 | 22322746 | N/A |
| PB19_depleted | Illumina HiSeqX | 2X151 | Smart-seq2 | 19418607 | N/A |
| PB21_depleted | Illumina HiSeqX | 2X151 | Smart-seq2 | 22467660 | N/A |
| PB3_undepleted | Illumina HiSeqX | 2X151 | Smart-seq2 | 58088942 | N/A |
| PB19_undepleted | Illumina HiSeqX | 2X151 | Smart-seq2 | 105936701 | N/A |
| PB21_undepleted | Illumina HiSeqX | 2X151 | Smart-seq2 | 63096050 | N/A |
| PB19_undepleted | ONT MinION | 9.4.1/LSK109 | R2C2 | 5313 | 93% |
| PB3_depleted_R1 | ONT MinION | 9.4.1/LSK109 | R2C2 | 390526 | 93% |
| PB3_depleted_R2 | ONT MinION | 9.4.1/LSK109 | R2C2 | 1691780 | 94% |
| PB19_depleted_R1 | ONT MinION | 9.4.1/LSK109 | R2C2 | 59097 | 93% |
| PB19_depleted_R2 | ONT MinION | 9.4.1/LSK109 | R2C2 | 866087 | 97.5% |
| PB21_depleted | ONT MinION | 9.4.1/LSK109 | R2C2 | 830952 | 92% |

**Table S2.**
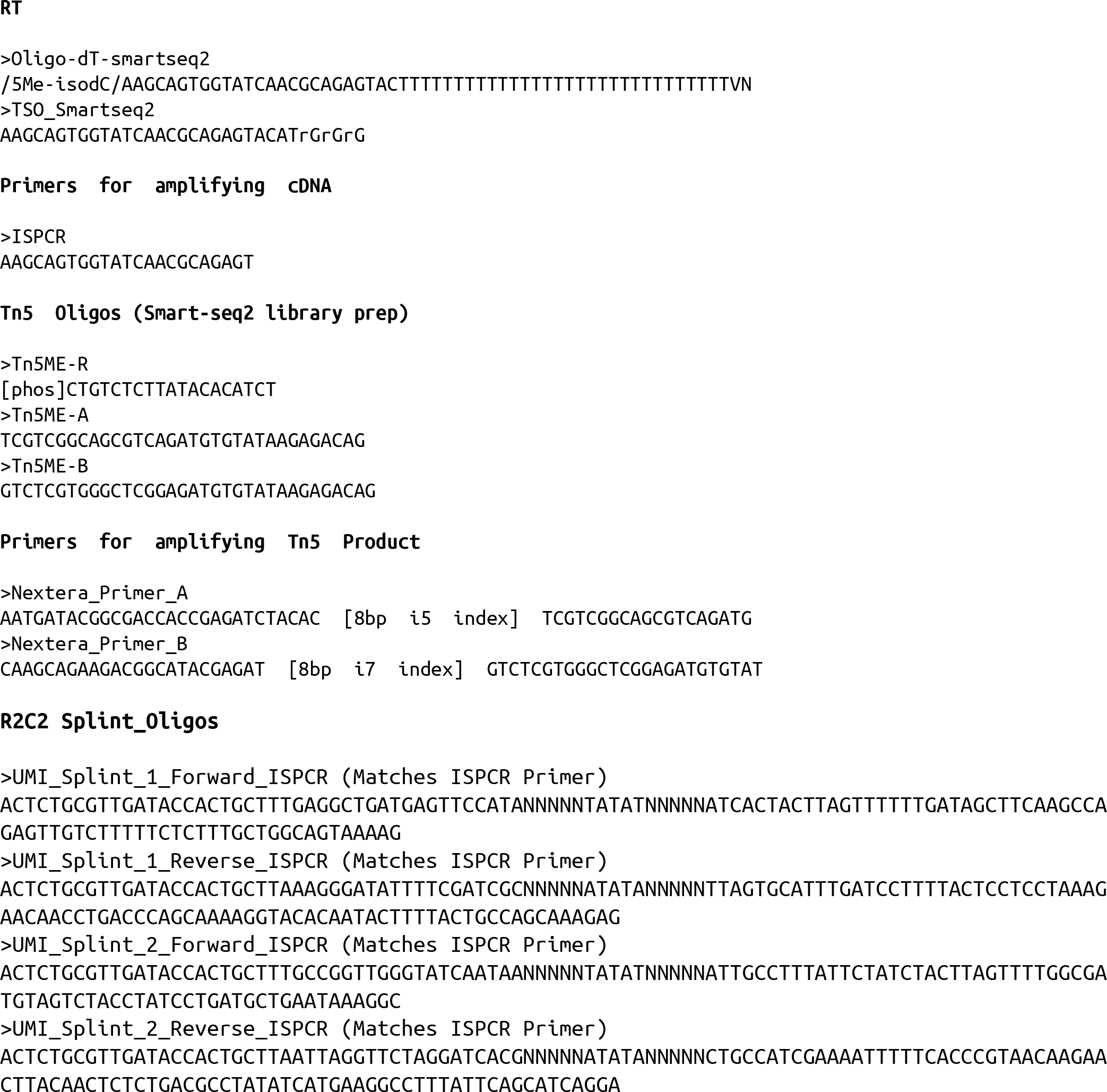
Oligos used in the manuscript. All oligos are shown 5’->3’ and were ordered from Integrated DNA Technologies (IDT). Lower case ‘r’ indicates RNA bases. Spaces in sequences are for visual emphasis only.

